# Polystyrene Microplastics Accelerate Antibiotic Resistance Evolution and Exacerbate Pathogenicity in *Acinetobacter Baumannii*

**DOI:** 10.64898/2026.08.27.747551

**Authors:** Muneer Yaqub, Namrata Bonde, Suman Tiwari, Ravali Arugonda, Tuhina Maity, William Cutts, Samuel Cornelius, Jon Sin, Nicole De Nisco, Ingrid Cornax, Brandon Kim, Nicholas Dillon

## Abstract

Microplastics are pervasive environmental contaminants and are increasingly detected in contexts relevant to human health, yet their effects on antimicrobial resistance, host–pathogen interactions, and infection outcomes remain poorly understood. Here, we show that exposure to polystyrene microplastics alters both antibiotic resistance evolution and pathogenic behavior in *Acinetobacter baumannii*, a leading cause of multidrug-resistant hospital-acquired infections. Using experimental evolution under antibiotic selection, we demonstrate that microplastic exposure accelerates resistance emergence across multiple antibiotic classes. Although microplastic exposure did not uniformly enhance biofilm formation, it modestly impaired macrophage-mediated bacterial clearance, suggesting broader effects on bacterial adaptation and host interaction. In vivo, microplastic-associated infection resulted in more severe disease, characterized by increased lung tissue damage and reduced survival in a murine model of *A. baumannii* pneumonia. Together, these findings identify microplastics as ecological modifiers of bacterial adaptation, linking widespread plastic pollution to enhanced antimicrobial resistance and worsened infectious disease outcomes.

## Introduction

Microplastics have emerged as pervasive environmental contaminants with growing implications for both ecosystem and human health. Commonly defined as plastic particles smaller than 5 mm, microplastics are now widely distributed across marine, freshwater, terrestrial, atmospheric, and clinical environments [1–3]. Their persistence, mobility, and increasing accumulation across biological systems have raised major concerns regarding their long-term ecological and biomedical consequences [4,5]. Beyond their environmental prevalence, microplastics are increasingly recognized as biologically active particles capable of interacting with microorganisms, host tissues, and environmental pollutants [6–8]. Their large surface area, hydrophobicity, and capacity to adsorb organic and inorganic compounds create unique physicochemical microenvironments that can influence microbial survival, adaptation, and interspecies interactions [9,10].

Growing evidence suggests that microplastics may function as ecological modifiers of microbial behavior. Environmental studies have shown that microplastics can serve as surfaces for microbial colonization and biofilm formation, generating “plastisphere” communities enriched with opportunistic pathogens and antibiotic resistance genes [11–14]. Exposure to microplastics has also been associated with altered bacterial stress responses, enhanced horizontal gene transfer, and reduced susceptibility to antimicrobial compounds in several bacterial species [15–18]. Recent studies further demonstrate that microplastics can directly enhance biofilm-associated antimicrobial resistance phenotypes and promote adaptive bacterial states under antibiotic stress [25–30]. In parallel, increasing attention has focused on the potential role of microplastics in shaping host immune responses and host–microbe interactions [10,12,18,31,32]. Despite these observations, the extent to which microplastic exposure influences bacterial adaptation and pathogenicity in clinically important pathogens remains poorly understood.

Among the pathogens of greatest concern in this context is *Acinetobacter baumannii*, an opportunistic Gram-negative bacterium recognized as a major global threat due to its extraordinary capacity for environmental persistence and multidrug resistance [33–36]. *A. baumannii* is a leading cause of hospital-acquired infections including ventilator-associated pneumonia, bloodstream infections, wound infections, and sepsis, particularly among critically ill and immunocompromised patients [33,34]. The increasing prevalence of carbapenem-resistant and extensively drug-resistant *A. baumannii* has severely limited treatment options, often leaving polymyxins and tetracycline derivatives among the few remaining effective therapies [37–40]. In addition to antimicrobial resistance, the pathogenic success of *A. baumannii* is strongly influenced by traits such as biofilm formation, resistance to oxidative and membrane stress, immune evasion, and survival within host cells [35,36]. These adaptive phenotypes enable *A. baumannii* to thrive across diverse environmental and host-associated niches.

Because microplastics can influence microbial aggregation, surface-associated growth, stress adaptation, and host immune function, they may substantially alter both antibiotic adaptation and virulence-associated behaviors in *A. baumannii*. However, despite increasing recognition of microplastics as ecological modifiers of microbial behavior, their impact on antibiotic adaptation and pathogenicity in *A. baumannii* has not been well defined.

In the present study, we used polystyrene microplastics as a representative microplastic type to investigate how microplastic exposure influences antibiotic resistance evolution, biofilm-associated phenotypes, host–pathogen interactions, and virulence in *A. baumannii* AB5075. Using experimental evolution approaches alongside in vitro macrophage assays and a murine pneumonia model, we examined how microplastic exposure shapes bacterial adaptation under antibiotic stress and affects infection outcomes during pulmonary disease. These findings provide insight into the emerging intersection between environmental microplastic contamination, antimicrobial resistance, and bacterial pathogenicity, highlighting the potential clinical relevance of microplastic exposure in shaping infection outcomes.

## Results

### Microplastic Exposure Accelerates the Evolution of Antibiotic Resistance in *A. baumannii*

To investigate whether polystyrene microplastics (MPs) influence the evolution of antibiotic resistance in *A. baumannii*, we performed parallel experimental evolution assays in the presence or absence of MPs under progressively increasing antibiotic selection pressure (Figure 1a). Four antibiotics were selected to span distinct mechanisms of action and clinically relevant resistance contexts: colistin, a last-resort polymyxin targeting the outer membrane; doxycycline and minocycline, tetracyclines that inhibit protein synthesis but differ in lipophilicity and intracellular accumulation; and meropenem, a carbapenem β-lactam targeting cell wall synthesis [37–40]. For each antibiotic, three independent AB5075 lineages were serially passaged daily under sub-inhibitory antibiotic exposure, with antibiotic concentrations increased in two-fold increments as populations adapted. Evolution experiments were continued until populations reached the maximum selection concentration of 1024 μg/ml or for a total of 14 days. Resistance trajectories were monitored for doxycycline, colistin, minocycline, and meropenem (Figure 1b–e).

**Figure 1.**
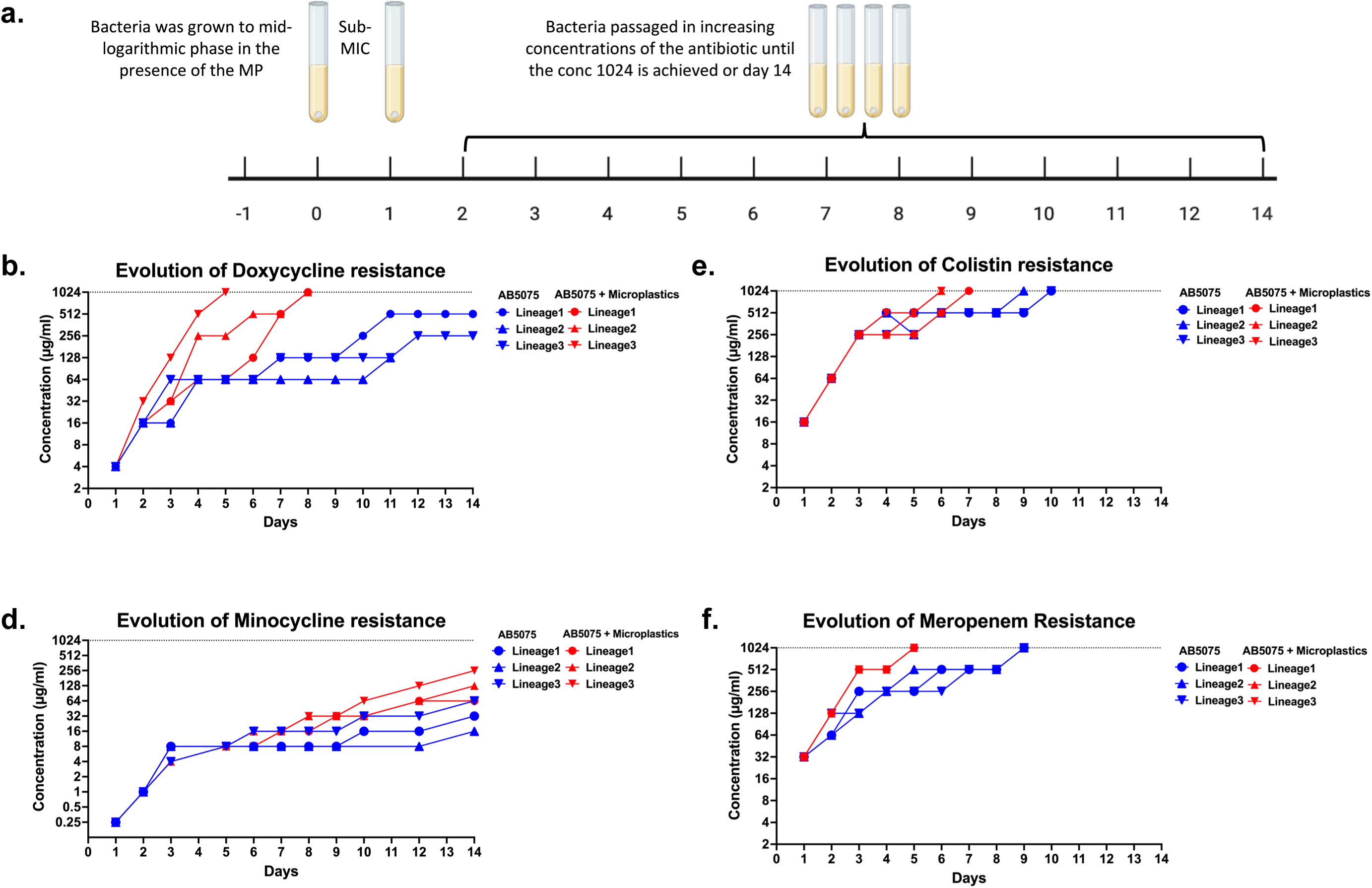
Experimental evolution of antibiotic resistance in the presence and absence of polystyrene microplastics. **(a)** Schematic of the experimental evolution protocol. Bacterial cultures were grown to mid-logarithmic phase under sub-inhibitory (sub-MIC) antibiotic conditions and serially passaged daily with and without microplastics in increasing antibiotic concentrations until a maximum of 1024 μg/ml was reached or up to 14 days. **(b–e)** Evolution of resistance across three independent lineages of *A. baumannii* AB5075 grown in the absence or presence of polystyrene microplastics. Resistance trajectories are shown for **(b)** doxycycline, **(c)** colistin, **(d)** minocycline, and **(e)** meropenem. Antibiotic concentrations (μg/ml) tolerated by each lineage are plotted over time (days). Each line represents an independent evolutionary lineage. Blue lines indicate AB5075, and red lines indicate AB5075 grown in the presence of polystyrene microplastics.

Across multiple antibiotic conditions, we found that MP exposure accelerated adaptive trajectories relative to MP-free populations. Under colistin selection, MP-exposed populations reached the maximum selection concentration substantially earlier than MP-free populations (mean ± SD: 6.3 ± 0.6 vs 9.7 ± 0.6 days) (Figure 1c; Extended Table 1). Although both conditions ultimately reached 1024 μg/ml by the end of the evolution period, MP-associated populations consistently advanced through the two-fold concentration series more rapidly.

A similar but more pronounced pattern emerged during doxycycline selection (Figure 1b). All MP-exposed populations reached 1024 μg/ml within the experimental timeframe (7.0 ± 1.7 days), whereas none of the MP-free populations achieved the maximum selection concentration by day 14 (Extended Table 1). Consistent with this divergence, MP-free populations reached substantially lower endpoint resistance levels than MP-associated populations by the conclusion of the experiment (Extended Table 2).

Under minocycline selection, neither condition reached 1024 μg/ml during the 14-day evolution period (Figure 1d). Nevertheless, MP-exposed populations achieved markedly higher final tolerated concentrations than MP-free populations, indicating accelerated adaptation despite incomplete convergence to the upper selection threshold (Extended Table 2).

Meropenem selection produced comparatively similar overall evolutionary trajectories between conditions, as both MP-free and MP-exposed populations ultimately reached 1024 μg/ml (Figure 1e). However, MP-associated populations achieved the maximum concentration substantially earlier than MP-free populations (5.0 ± 0.0 vs 9.0 ± 0.0 days), demonstrating that MPs also accelerated adaptation dynamics under carbapenem selection (Extended Table 1).

Together, these findings demonstrate that polystyrene MPs can accelerate the evolution of antibiotic resistance in *A. baumannii* across multiple antibiotic classes. While the magnitude of the effect varied between antibiotics, MP-associated populations generally progressed through increasing antibiotic concentrations more rapidly and achieved higher endpoint resistance levels than populations evolved in the absence of MPs.

### Microplastic exposure does not universally enhance biofilm formation of *A. baumannii*

Because biofilm formation can contribute to antibiotic tolerance and adaptive survival under stress, we next examined whether the accelerated resistance evolution observed in MP-exposed populations was associated with altered biofilm formation. Biofilm production was quantified across independently evolved *A. baumannii* AB5075 populations generated under doxycycline, colistin, minocycline, and meropenem selection in the presence or absence of polystyrene microplastics (MPs) using standard crystal violet biofilm assays (Figure 2a–d).

**Figure 2.**
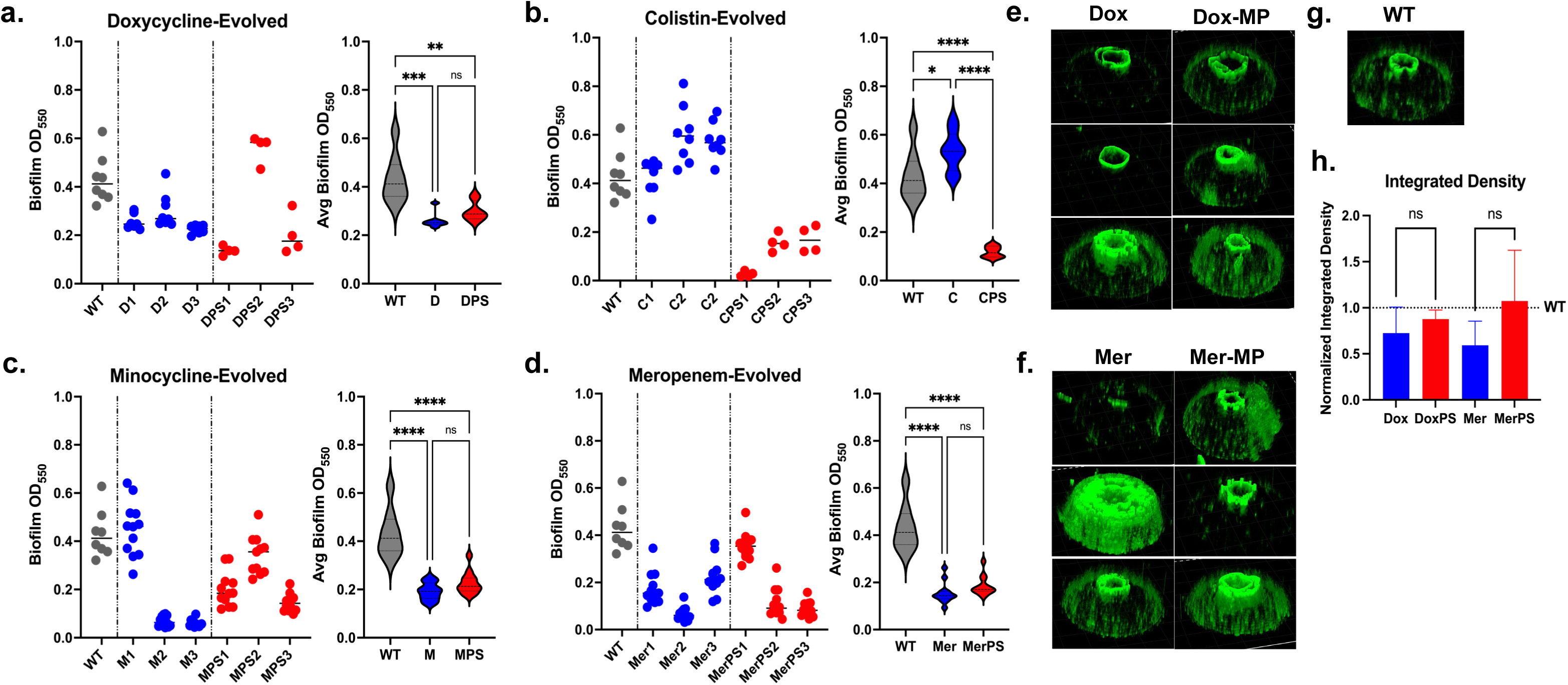
Polystyrene microplastics modulate biofilm formation in antibiotic-evolved *A. baumannii* AB5075 populations. (a–d) Biofilm formation (OD₅₅₀) of evolved lineages following selection with **(a)** doxycycline, **(b)** colistin**, (c)** minocycline, and **(d)** meropenem. Individual data points represent independent evolved populations grown in the absence (blue) or presence (red) of polystyrene microplastics, with wild-type (WT, gray) shown for comparison. Corresponding violin plots summarize average biofilm production across groups. **(e–f)** Representative 3D confocal images of biofilms formed under **(e)** doxycycline and **(f)** meropenem selection, in the absence or presence of polystyrene microplastics (MP). **(g)** Representative biofilm architecture of the wild-type strain. **(h)** Quantification of biofilm biomass based on normalized integrated fluorescence intensity across conditions. Statistical significance is indicated where applicable (ns, not significant; *p < 0.05; **p < 0.01; ***p < 0.001; ****p < 0.0001).

Across most antibiotic conditions, MP exposure did not produce a consistent global increase in biofilm formation. Under doxycycline selection, biofilm production was broadly comparable between MP-exposed and MP-free populations despite variability among individual evolved lineages (Figure 2a). Similarly, populations evolved under meropenem selection displayed largely overlapping biofilm phenotypes irrespective of MP exposure (Figure 2d). Minocycline-selected populations exhibited modest variation between conditions, although this effect was not consistently observed across all lineages (Figure 2c). In contrast, the clearest separation emerged under colistin selection, where MP-exposed populations demonstrated elevated biofilm formation relative to MP-free populations (Figure 2b).

To further assess structural differences in biofilm organization, representative biofilms formed under doxycycline and meropenem selection were visualized by confocal microscopy (Figure 2e,f). These analyses revealed heterogeneous three-dimensional biofilm architectures across evolved populations, including differences in thickness, topology, and spatial organization. However, quantitative analysis of normalized integrated fluorescence intensity showed no consistent increase in total biomass associated with MP exposure across conditions (Figure 2h). Representative imaging of the wild-type AB5075 strain provided a baseline reference for comparison with evolved populations (Figure 2g).

Importantly, biofilm phenotypes did not parallel the resistance trajectories observed during experimental evolution. MP exposure accelerated resistance evolution across all antibiotics tested, including doxycycline and meropenem, despite the absence of corresponding increases in biofilm formation under those conditions. Likewise, although enhanced biofilm production was observed under colistin selection, this phenotype was not universally conserved across all antibiotic environments. Together, these findings indicate that enhanced biofilm formation is unlikely to represent a general mechanism underlying MP-associated acceleration of antibiotic resistance evolution in *A. baumannii*. Rather, the effects of MPs on biofilm formation appear to be antibiotic-dependent, with a more pronounced biofilm-associated phenotype emerging specifically under colistin selection.

### Polystyrene microplastics modestly impair macrophage-mediated clearance of *A. baumannii*

To determine whether polystyrene microplastics (MPs) influence host–pathogen interactions during *A. baumannii* infection, we evaluated bacterial survival and macrophage viability during infection of THP-1–derived macrophages under conditions with or without MP exposure (Figure 3). Macrophage killing assays were performed over a 120-minute infection period to assess intracellular bacterial survival dynamics and host cell responses.

**Figure 3.**
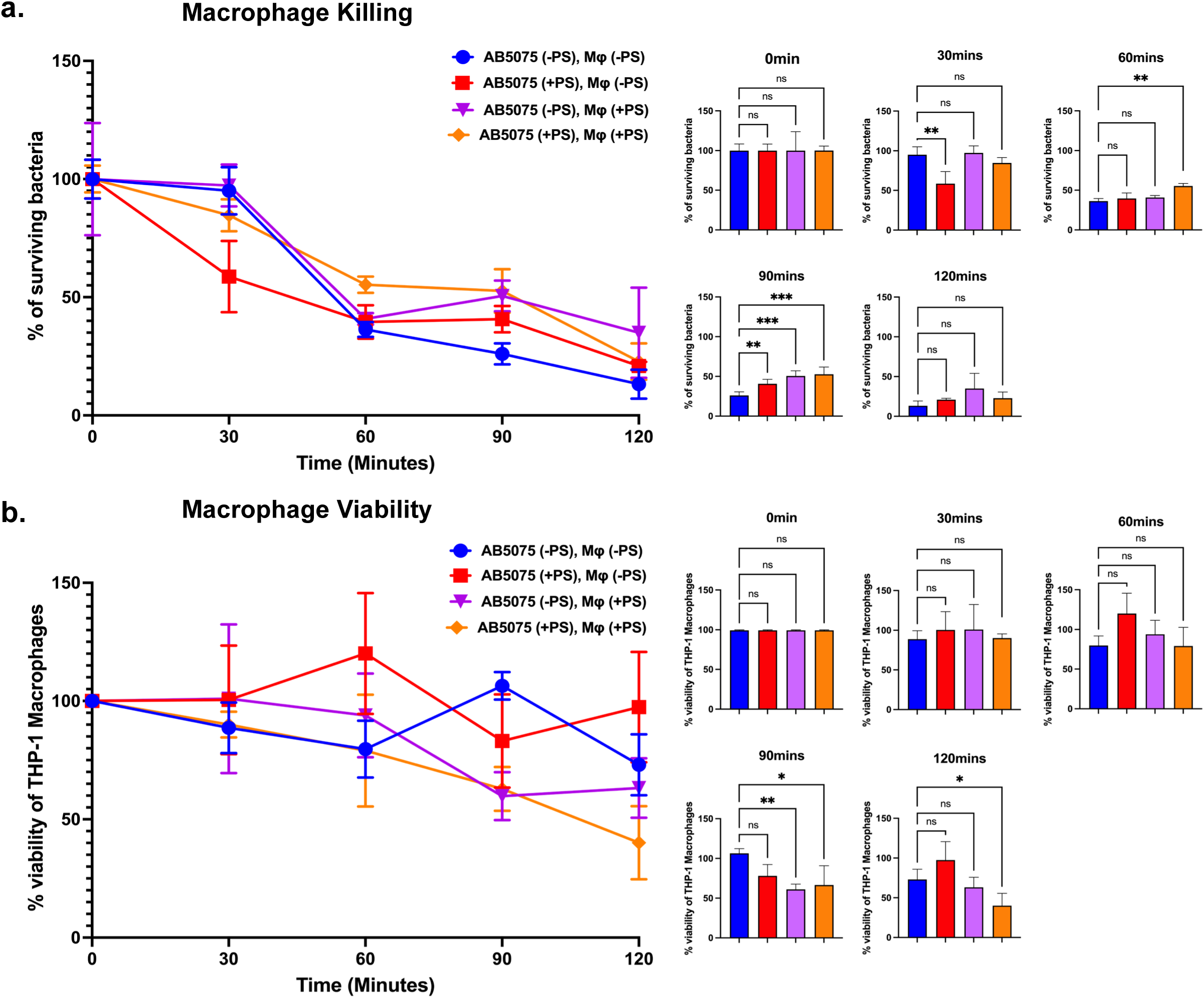
Polystyrene microplastics alter macrophage–*A. baumannii* AB5075 interactions. **(a)** Macrophage killing assay showing the percentage of surviving *A. baumannii* AB5075 over time (0–120 min) following infection of THP-1–derived macrophages (Mφ). Bacteria and macrophages were exposed to conditions with or without polystyrene microplastics (PS), as indicated. Line graphs depict bacterial survival kinetics, with corresponding bar plots showing time-point comparisons. **(b)** Macrophage viability assay showing the percentage viability of THP-1 macrophages over time (0–120 min) under the same conditions. Line graphs represent temporal changes in host cell viability, with accompanying bar plots for individual time points. Data are presented as mean ± standard deviation. Statistical significance is indicated where applicable (ns, not significant; *p < 0.05; **p < 0.01; ***p < 0.001).

Macrophages efficiently reduced bacterial burden over time across all conditions, reflecting progressive macrophage-mediated clearance of *A. baumannii* AB5075 (Figure 3a). However, exposure to MPs altered the kinetics of bacterial survival during infection. Although early time points showed broadly similar survival profiles, MP-associated conditions generally maintained modestly higher bacterial survival at later stages of the assay compared with

MP-free conditions. This effect was observed whether MPs were present during bacterial exposure, macrophage exposure, or both, suggesting that MPs can influence host–pathogen interactions through effects on either the bacterium, the host cell, or both simultaneously. Despite these trends, differences between conditions were relatively moderate and did not produce complete impairment of macrophage killing capacity.

To assess whether altered bacterial survival reflected overt macrophage toxicity, macrophage viability was monitored throughout the infection period (Figure 3b). Across all conditions, THP-1 macrophage viability remained relatively stable over time, with only modest fluctuations between groups. MP exposure alone did not produce substantial reductions in host cell viability, and infection-associated decreases in viability were limited overall. These findings indicate that the altered bacterial survival dynamics observed in MP-associated conditions were not primarily attributable to widespread macrophage death or gross cytotoxicity.

Together, these results suggest that polystyrene MPs modestly impair macrophage-mediated clearance of *A. baumannii* without causing major loss of macrophage viability. The dissociation between bacterial survival and host cell death further suggests the possibility that MPs subtly modulate host–pathogen interactions and innate immune function during *A. baumannii* infection.

### Polystyrene microplastics enhance *A. baumannii* virulence in a murine pneumonia model

To determine whether the altered host–pathogen interactions observed in vitro translated to infection outcomes in vivo, we evaluated the impact of polystyrene microplastics (MPs) on *A. baumannii* virulence using a murine pneumonia model (Figure 4a). Female and male C57BL/6J mice were infected intratracheally with *A. baumannii* AB5075 in the presence or absence of MPs, and survival was monitored over a 6-day period.

**Figure 4.**
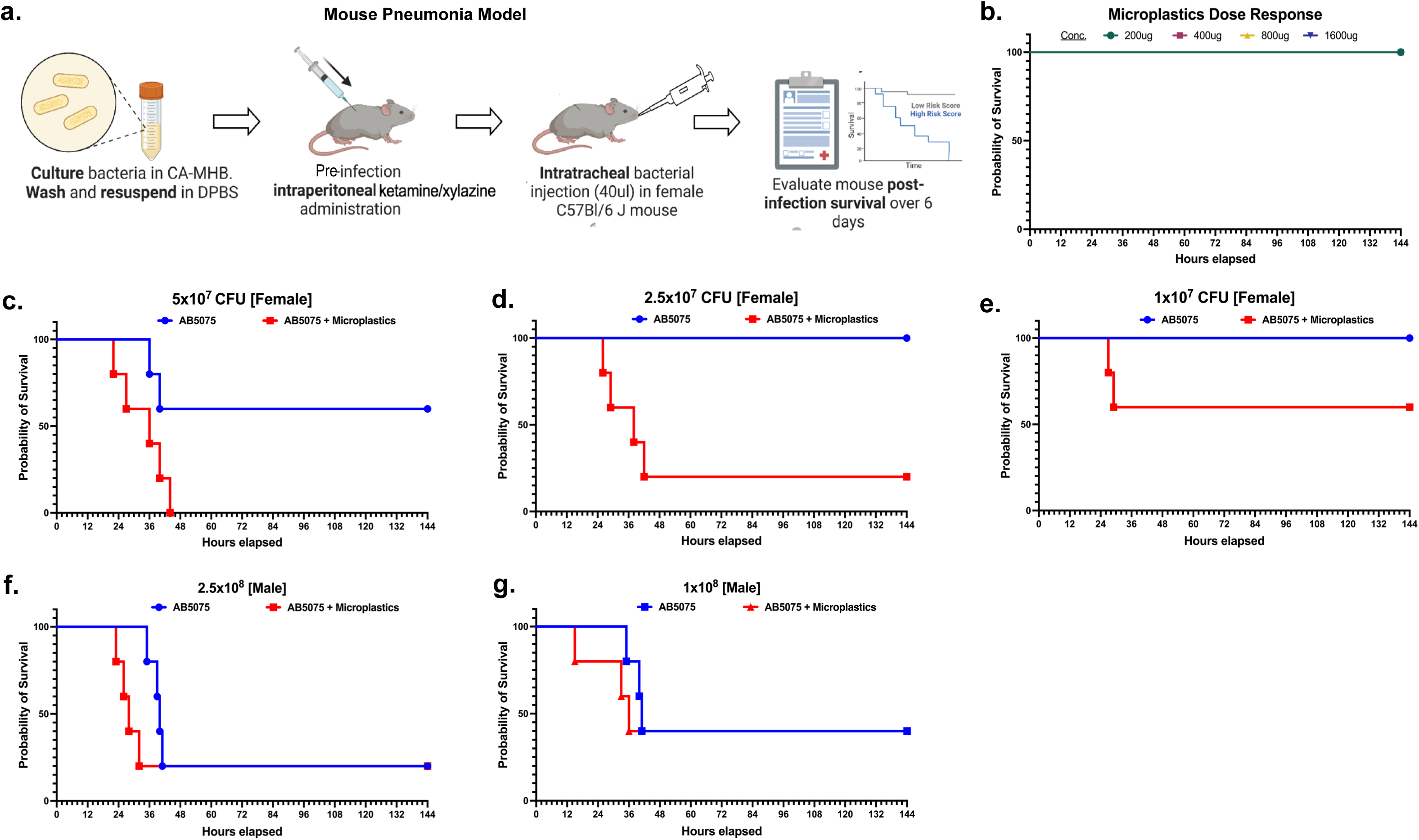
Polystyrene microplastics enhance virulence in a murine pneumonia model of *A. baumannii* AB5075. **(a)** Schematic of the mouse pneumonia infection model. Bacterial cultures were prepared, washed, and resuspended prior to intratracheal inoculation in anesthetized mice, with or without polystyrene microplastics. Survival was monitored over a 6-day period. **(b)** Survival analysis of mice exposed to increasing doses of polystyrene microplastics alone, showing no significant effect on host survival across tested concentrations. **(c–e)** Kaplan–Meier survival curves of mice infected with *A. baumannii* AB5075 in the absence (blue) or presence (red) of polystyrene microplastics at varying inoculum sizes: (c) 5×10⁷ CFU, (d) 2.5×10⁷ CFU, and **(e)** 1×10⁷ CFU. **(f–g)** Survival analysis in male mice infected with higher bacterial inocula: **(f)** 2.5×10⁸ CFU and **(g)** 1×10⁸ CFU, with or without polystyrene microplastics. Survival is presented as Kaplan–Meier curves over time (hours post-infection). Blue lines indicate AB5075 alone, and red lines indicate AB5075 in the presence of polystyrene microplastics.

To first assess whether MPs alone produced overt toxicity under the experimental conditions, mice were exposed to increasing MP doses in the absence of bacterial infection. Across all tested concentrations, MP exposure alone did not significantly affect host survival, with all animals remaining viable throughout the observation period (Figure 4b). These findings indicate that the mortality phenotypes observed during infection were not attributable to acute MP toxicity alone.

In contrast, co-exposure to MPs markedly altered infection outcomes during *A. baumannii* pneumonia. At an inoculum of 5 × 10⁷ CFU, mice infected in the presence of MPs exhibited substantially accelerated mortality compared with animals infected with bacteria alone (Figure 4c). While AB5075 infection in the absence of MPs resulted in delayed and incomplete mortality over the experimental period, MP-associated infection produced rapid and near-complete mortality within a substantially shorter timeframe.

A similar pattern was observed at lower inocula. At 2.5 × 10⁷ CFU, all mice infected with AB5075 alone survived the duration of the experiment, whereas co-exposure with MPs resulted in pronounced mortality (Figure 4d). Likewise, at 1 × 10⁷ CFU, MP-associated infection reduced host survival relative to infection with bacteria alone, despite the lower bacterial burden (Figure 4e). These findings demonstrate that MPs enhance disease severity across a range of infectious doses, including conditions that are otherwise minimally lethal in the absence of MPs.

To further examine whether this phenotype extended across sex and higher infectious burdens, male mice were infected with elevated inocula of AB5075. At 2.5 × 10⁸ CFU, both MP-free and MP-associated infections ultimately resulted in complete mortality; however, MP exposure accelerated the kinetics of death relative to infection alone (Figure 4f). Similarly, at 1 × 10⁸ CFU, MP-associated infection produced reduced survival and earlier mortality compared with AB5075 infection in the absence of MPs (Figure 4g).

Together, these findings demonstrate that polystyrene MPs significantly exacerbate *A. baumannii* virulence during pulmonary infection.

### Microplastic Exposure Augments Lung Tissue Damage during *A. baumannii* Infection

Because microplastic exposure was associated with reduced host survival during *A. baumannii* lung infection, we next assessed whether this was accompanied by increased lung tissue damage. To address this, lung sections were collected from infected mice and analyzed by histopathology, with tissue injury scored using a semi-quantitative scale assessing inflammation and tissue injury.

In uninfected control mice, lung architecture was preserved, with thin alveolar septa and minimal inflammatory infiltrates (Figure 5a). Exposure to polystyrene microplastics alone resulted in minimal accumulation of inflammatory neutrophilic infiltrates around terminal airways (Figure 5b), consistent with the absence of overt disease or mortality observed in microplastic-only controls. Infection with AB5075 in the absence of microplastics resulted in histologic changes consistent with bacterial pneumonia, including neutrophilic inflammation and multifocal hemorrhage (Figure 5c). In the combination group, the overall severity of neutrophilic inflammation and hemorrhage was comparable to that observed with AB5075 alone. However, peripheral proteinaceous edema and necrotic cellular debris were observed within the subpleural alveolar spaces, consistent with disruption of alveolar wall integrity (Figure 5d). Polystyrene exposure alone was associated with minimal histopathological changes.

**Figure 5.**
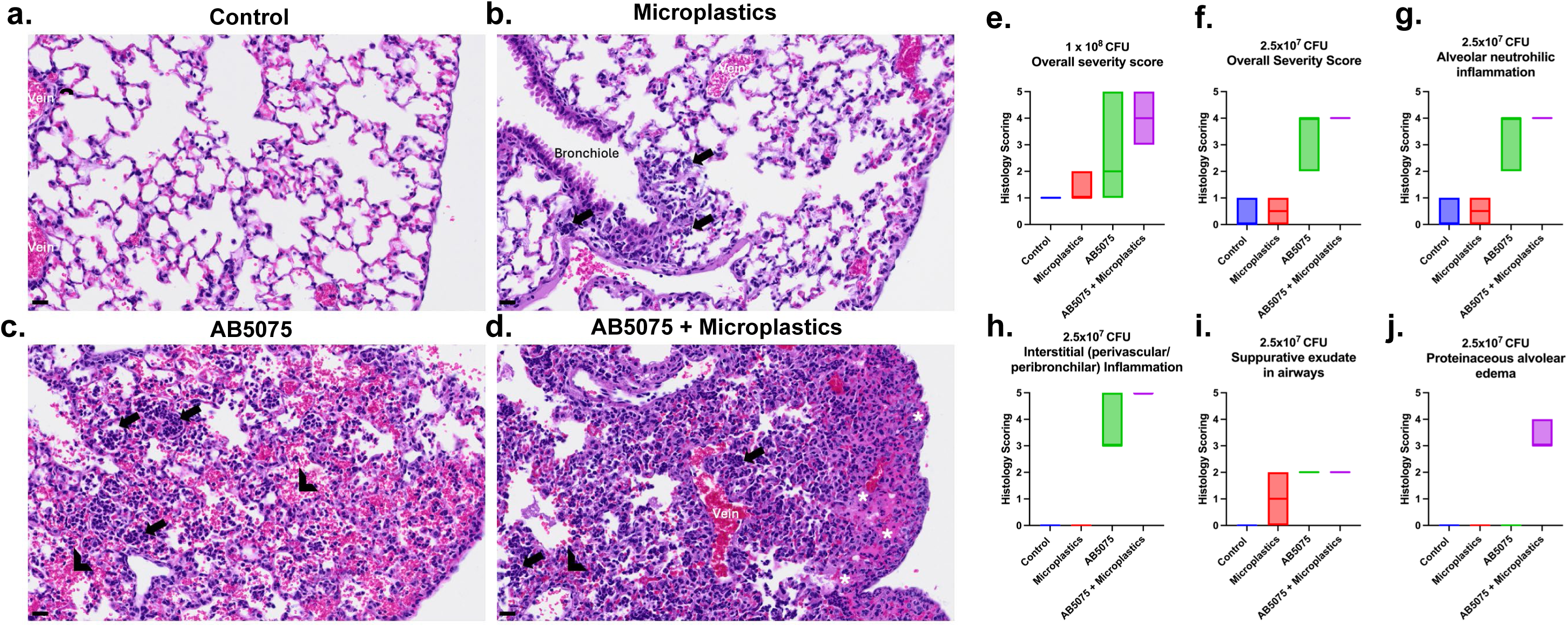
Representative histopathology and semiquantitative scoring of lung injury following microplastic exposure and *A. baumannii* AB5075 infection. **(a-d)** Representative microscopic images of lungs from each treatment group. **(a)** uninfected control showing normal air spaces and blood contained within blood vessels (veins). **(b)** polystyrene only. Small foci of neutrophilic inflammation (arrows) present at terminal airways. Air spaces are otherwise normal. **(c)** AB5705 infection. Multiple aggregates of degenerate neutrophils (arrows) and hemorrhage (chevrons) present air spaces. **(d)** combination polystyrene and AB5705 infection demonstrating presence of proteinaceous edema fluid and necrotic debris within air spaces (asterisks) in addition to neutrophilic inflammation (arrows) and hemorrhage (chevrons). Hematoxylin and eosin-stained sections, scale bar = 20 µm. **(e–j)** Semiquantitative histopathology scoring across treatment groups. **(e)** Overall severity score at 1×10^8^ CFU challenge. **(f)** Overall severity score at 2.5×10^7^ CFU challenge. **(g)** Alveolar neutrophilic inflammation at 2.5×10^7^ CFU. **(h)** Interstitial (perivascular/peribronchiolar) inflammation at 2.5×10^7^ CFU. **(i)** Suppurative exudate within airways at 2.5 × 10^7 CFU. **(j)** Proteinaceous alveolar edema at 2.5 × 10^7 CFU. Each point represents an individual animal; boxes indicate score distribution across groups.

Together, these observations suggest that polystyrene exposure may alter the pattern of lung pathology during *A. baumannii* infection and provide histological context for the reduced host survival observed following combined exposure.

### 3.4 Discussion

Our findings demonstrate that polystyrene MPs can substantially alter the adaptive and pathogenic behavior of *A. baumannii* across multiple biological scales, including antibiotic resistance evolution, host–pathogen interactions, and in vivo virulence. Using experimental evolution, macrophage infection assays, and murine pneumonia models, we show that MP exposure accelerates resistance acquisition under diverse antibiotic selection pressures while simultaneously exacerbating disease severity during pulmonary infection. Together, these results support the growing view that microplastics are not merely passive environmental contaminants, but biologically active ecological modifiers capable of reshaping bacterial adaptation and infection outcomes.

One of the most striking observations in this study was the consistent acceleration of resistance evolution in MP-associated populations. Across several antibiotic classes, including tetracyclines, polymyxins, and carbapenems, MP-exposed lineages progressed through increasing antibiotic concentrations more rapidly than populations evolved in the absence of MPs. These findings align with accumulating evidence suggesting that microplastics can generate physicochemical microenvironments that facilitate bacterial adaptation under stress [9–30,41,42]. MPs have been proposed to promote bacterial aggregation, alter nutrient availability, adsorb antimicrobial compounds, and induce stress-response pathways that collectively increase adaptive potential. Recent work further demonstrates that microplastics can directly enhance biofilm-associated antimicrobial resistance and promote persistent adaptive phenotypes even after particle removal [25]. In our system, however, the effects of MPs were not uniform across all antibiotics, indicating that the interaction between MPs and antibiotic stress is likely context dependent and influenced by the underlying mechanism of antimicrobial action.

Importantly, enhanced biofilm formation did not emerge as a universal explanation for the accelerated resistance trajectories observed in MP-associated populations. Although increased biofilm production was evident under colistin selection, most antibiotic conditions showed only modest or inconsistent differences between MP-exposed and MP-free populations. Confocal imaging similarly failed to reveal broad increases in biofilm biomass associated with MPs. These findings suggest that while biofilm-associated phenotypes may contribute under specific conditions, particularly during polymyxin exposure, they are unlikely to represent the primary mechanism underlying MP-associated acceleration of resistance evolution. Instead, the present findings raise the possibility that MPs may influence bacterial adaptation through biofilm-independent mechanisms. Potential areas for future investigation include membrane stress responses, oxidative stress adaptation, metabolic restructuring, and altered antibiotic diffusion dynamics within particle-associated microenvironments.

The pronounced biofilm phenotype observed specifically under colistin selection is nevertheless notable. Colistin exerts bactericidal activity through disruption of the Gram-negative outer membrane [37], and adaptive responses to polymyxin stress frequently involve extensive remodeling of surface architecture, membrane charge, and extracellular matrix production. It is therefore plausible that MPs interact more strongly with cellular pathways linked to envelope stress and surface adaptation during colistin exposure than during other antibiotic conditions. Whether this reflects altered particle–membrane interactions, enhanced extracellular polymeric substance production, or selective enrichment of surface-associated subpopulations remains to be determined.

Beyond resistance evolution, our results further demonstrate that MPs can alter host–pathogen interactions during infection. In macrophage infection assays, MP-associated conditions produced modest but reproducible impairment of macrophage-mediated bacterial clearance without causing substantial reductions in macrophage viability. This dissociation between bacterial persistence and overt cytotoxicity suggests that MPs subtly modulate innate immune function rather than simply inducing nonspecific host cell death. Prior studies have shown that microplastics can influence macrophage inflammatory signaling, phagocytic capacity, oxidative burst responses, and cytokine production [5,10,12,18]. Our findings extend these observations to *A. baumannii*, indicating that MPs may create conditions that favor bacterial persistence within the host even in the absence of severe immune cell toxicity.

Consistent with the in vitro macrophage phenotypes, MP exposure dramatically worsened infection outcomes in vivo. Across multiple inoculum sizes and in both female and male mice, co-exposure to MPs accelerated mortality and increased disease severity during *A. baumannii* pneumonia. Importantly, MPs alone did not cause measurable mortality, indicating that the observed phenotypes reflected enhancement of bacterial pathogenicity rather than direct toxic effects of particle exposure alone. Histopathologic analysis further demonstrated that MP-associated infection was accompanied by increased pulmonary tissue damage, including edema formation and loss of alveolar integrity. Together, these findings suggest that MPs potentiate pulmonary disease through combined effects on bacterial adaptation, host immune interactions, and tissue injury responses.

The mechanisms linking MPs to enhanced virulence remain incompletely understood. One possibility is that MPs alter bacterial surface properties or stress physiology in ways that increase resistance to host clearance mechanisms. Alternatively, MPs may directly modulate pulmonary immune responses, impairing early containment of infection and promoting excessive inflammatory damage. MPs can also adsorb proteins, lipids, and other biologically active molecules, potentially creating localized microenvironments that influence both bacterial and host cell behavior during infection. The observation that MPs enhanced virulence even at relatively low infectious doses raises the possibility that environmentally relevant particle exposure could meaningfully influence infection outcomes under clinical conditions.

Several limitations should also be considered. The present study focused specifically on polystyrene MPs, whereas environmental microplastic exposure consists of highly heterogeneous particles varying in polymer composition, size, shape, charge, and chemical aging state. Different classes of MPs may therefore exert distinct biological effects. In addition, our experimental systems modeled acute exposure conditions, which may not fully recapitulate chronic environmental or occupational microplastic exposure experienced by humans. The molecular mechanisms linking MP exposure to accelerated resistance evolution and enhanced virulence also remain unresolved and warrant future mechanistic investigation.

Overall, our study demonstrates that polystyrene MPs can simultaneously accelerate antibiotic resistance evolution and exacerbate bacterial virulence in *A. baumannii*. These findings highlight microplastics as emerging environmental factors capable of influencing both antimicrobial resistance dynamics and infectious disease severity. As environmental MP contamination continues to increase globally, understanding how these particles shape microbial adaptation and host susceptibility may become increasingly important for both environmental microbiology and infectious disease research.

## Materials and Methods

### Experimental Evolution

*A. baumannii* AB5075 was used for all experimental evolution assays [25]. Cultures were maintained in cation-adjusted Mueller–Hinton broth (CAMHB) at 37 °C with shaking. For experimental evolution, replicate populations were serially passaged under stepwise increasing concentrations of colistin, doxycycline, or minocycline in the presence or absence of polystyrene (PS) microplastics (500 µm; Polysciences). Initial antibiotic concentrations were set at 0.25× MIC of the ancestral strain, doubling at each passage until resistance exceeded clinical breakpoints. Daily transfers were performed into fresh medium with antibiotics and PS (1 mg mL⁻¹), with controls lacking PS. Cultures were maintained for 10–14 days or until high-level resistance was achieved.

To compare resistance evolution dynamics between conditions, two complementary metrics were quantified from experimental evolution trajectories: (i) the time required for populations to reach the maximum selection concentration (1024 μg/ml), and (ii) the highest tolerated antibiotic concentration achieved by day 14. Because antibiotic concentrations were increased in two-fold increments during serial passaging, resistance progression was evaluated as advancement through discrete concentration steps rather than continuous MIC shifts. For each condition, values from three independent evolutionary lineages were summarized as mean ± standard deviation (SD). Populations that did not attain 1024 μg/ml within the experimental period were designated as “not reached within 14 days.”

### MIC Determination

Minimum inhibitory concentrations (MICs) were determined by broth microdilution according to CLSI guidelines [44]. Overnight cultures in the presence and absence of polystyrene were subcultured to mid-log phase, diluted to OD₆₀₀ = 0.002 in CAMHB, and inoculated into 96-well plates containing two-fold serial dilutions of antibiotics (colistin, doxycycline, minocycline, meropenem). Plates were incubated for 18–20 h at 37 °C, and MIC values were defined as the lowest concentration with no visible growth. Evolved lineages were compared with the parental strain, and collateral sensitivity or cross-resistance profiles were calculated as fold-changes in MIC relative to the ancestor.

### Biofilm Assay

Biofilm biomass was quantified using a crystal violet staining assay [45]. Mid-log cultures were normalized to OD₆₀₀ = 0.05 in CAMHB and inoculated into sterile 96-well plates (200 µL per well). Plates were incubated statically at 37 °C for 24 h. Non-adherent cells were removed by washing with distilled water, and wells were stained with 0.1% crystal violet for 15 min. Excess dye was removed, and biofilms were solubilized with 30% acetic acid. Absorbance was measured at 550 nm using a plate reader. Each assay included technical triplicates and three biological replicates.

### Macrophage Infection Assay

Intracellular survival of *A. baumannii* was assessed using a THP-1 macrophage infection assay [46]. THP-1 macrophages were cultured in DMEM supplemented with 10% fetal bovine serum at 37 °C with 5% CO₂. Cells were seeded at 2 × 10⁵ per well in 24-well plates and infected with mid-log *A. baumannii* AB5075 at a multiplicity of infection (MOI) of 10:1. After 30 min of infection, extracellular bacteria were removed by washing and replaced with media containing gentamicin (100 µg mL⁻¹). Intracellular bacteria were quantified at 30, 60, and 90 min post-infection by lysing macrophages with 0.1% Triton X-100, plating serial dilutions on LB agar, and enumerating colony-forming units (CFU). For co-exposure conditions, PS particles (1 mg mL⁻¹) were pre-incubated with either bacteria or macrophages for 2 h prior to infection.

### Murine Pneumonia Model

Murine pneumonia infections were performed using a previously established intratracheal infection model [47]. All animal experiments were conducted in accordance with institutional guidelines and approved by the Institutional Animal Care and Use Committee (IACUC). Female and male C57BL/6 mice (7–9 weeks old) were anesthetized with ketamine/xylazine and inoculated intratracheally with *A. baumannii* AB5075 at doses ranging from 5 × 10⁶ to 2.5 × 10⁸ CFU, suspended in 40 µL PBS with or without PS particles (400 µg). Survival was monitored for 6 days. Control groups received either PS alone (200–1600 µg) or bacteria alone.

### Histopathological Analysis

Lung histopathology was assessed using standard hematoxylin and eosin (H&E) staining procedures [48]. At 24 h post-infection, lungs were perfused with pbs and fixed in 4% paraformaldehyde overnight. Tissues were embedded in paraffin, sectioned at 5 µm, and stained with H&E. Slides were examined under a light microscope by a board-certified veterinary pathologist (IC), and representative images were captured at 200× magnification. Histological features were assessed for alveolar integrity, inflammation, and hemorrhage. Images shown are representative of at least three animals per group.

## Acknowledgement

This work was supported in part by seed funding from The University of Texas at Dallas and by funding from the National institutes of Health (grants AI168159 and GM143053 to JMB). We thank the UT Dallas Animal Resource Center for assistance with animal husbandry and Dr. Xintong Dong for help with the in vivo work.

---

**Extended Table 1.**
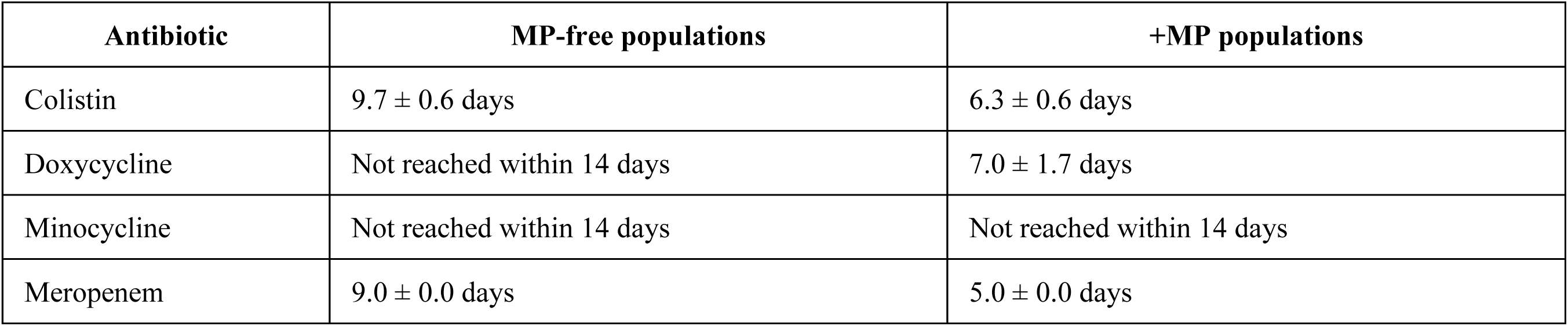
Time required to reach the maximum selection concentration (1024 μg/ml) Time required for independently evolved *A. baumannii* AB5075 populations to reach the maximum selection concentration during experimental evolution in the presence or absence of polystyrene microplastics (MPs). Experimental evolution was performed under progressively increasing antibiotic concentrations using daily serial passaging in two-fold concentration increments. Values represent the mean ± standard deviation (SD) number of days required for three independent lineages to reach the maximum selection concentration of 1024 μg/ml. “Not reached within 14 days” indicates that populations did not attain 1024 μg/ml within the 14-day experimental period.

**Extended Table 2.**
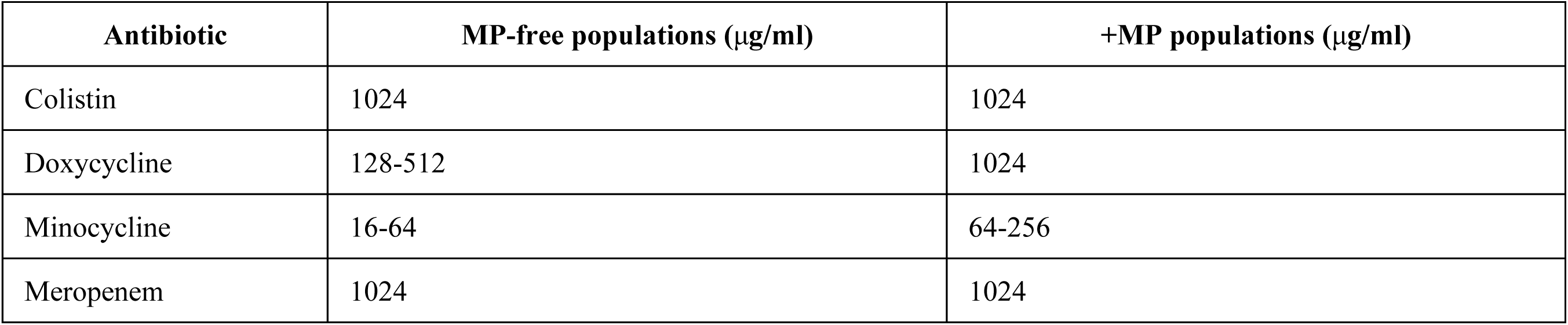
Endpoint antibiotic concentrations after 14 days of experimental evolution. Final antibiotic selection concentrations attained by evolved *A. baumannii* AB5075 populations after 14 days of experimental evolution in the presence or absence of polystyrene microplastics (MPs). Experimental evolution was performed under progressively increasing antibiotic selection pressure using daily serial passaging in two-fold concentration increments. Values represent the highest antibiotic concentration (μg/ml) reached by each independently evolved lineage at the end of the 14-day evolution experiment.

## References

1. Luo, Q., Tan, H., Ye, M., Jho, E. H., Wang, P., Iqbal, B., Zhao, X., Shi, H., Lu, H., & Li, G. (2025). Microplastics as an emerging threat to human health: An overview of potential health impacts. Journal of Environmental Management, 387, 125915. 10.1016/j.jenvman.2025.125915

2. Imran, M. (2025). Microplastic contamination: A rising environmental crisis with potential oncogenic implications. Cureus, 17(6), e85191. 10.7759/cureus.85191

3. Thompson, R. C., Courtene-Jones, W., Boucher, J., Pahl, S., Raubenheimer, K., & Koelmans, A. A. (2024). Twenty years of microplastic pollution research—What have we learned? Science, 386(6720), eadl2746. 10.1126/science.adl2746

4. Roslan, N. S., Lee, Y. Y., Ibrahim, Y. S., Tuan Anuar, S., Yusof, K. M. K. K., Lai, L. A., & Brentnall, T. (2024). Detection of microplastics in human tissues and organs: A scoping review. Journal of Global Health, 14, 04179. 10.7189/jogh.14.04179

5. Lee, Y., Cho, J., Sohn, J., & Kim, C. (2023). Health effects of microplastic exposures: Current issues and perspectives in South Korea. Yonsei Medical Journal, 64(5), 301–308. 10.3349/ymj.2023.0048

6. Anuar, S. T., Abdullah, N. S., Yahya, N. K. E. M., Chin, T. T., Yusof, K. M. K. K., Mohamad, Y., Azmi, A. A., Jaafar, M., Mohamad, N., Khalik, W. M. A. W. M., & Ibrahim, Y. S. (2023). A multidimensional approach for microplastics monitoring in two major tropical river basins, Malaysia. Environmental Research, 227, 115717. 10.1016/j.envres.2023.115717

7. Daud, A., & Astuti, R. D. P. (2021). Detection of exposure to microplastics in humans: A systematic review. Open Access Macedonian Journal of Medical Sciences, 9(F), 275– 280.

8. Danopoulos, E., Twiddy, M., & Rotchell, J. M. (2020). Microplastic contamination of drinking water: A systematic review. PLOS ONE, 15(7), e0236838. 10.1371/journal.pone.0236838

9. Nath, J., De, J., Sur, S., & Banerjee, P. (2023). Interaction of microbes with microplastics and nanoplastics in agroecosystems—Impact on antimicrobial resistance. Pathogens, 12(7), 888. 10.3390/pathogens12070888

10. Yee, M. S., Hii, L. W., Looi, C. K., Lim, W. M., Wong, S. F., Kok, Y. Y., Tan, B. K., Wong, C. Y., & Leong, C. O. (2021). Impact of microplastics and nanoplastics on human health. Nanomaterials, 11(2), 496. 10.3390/nano11020496

11. Microplastic research must consider microbes. (2025). Nature Microbiology, 10, 603. 10.1038/s41564-025-01960-6

12. Pacher-Deutsch, C., Schweighofer, N., Hanemaaijer, M., Marut, W., Žukauskaitė, K., Horvath, A., & Stadlbauer, V. (2025). The microplastic crisis: Role of bacteria in fighting microplastic effects in the digestive system. Environmental Pollution, 366, 125437. 10.1016/j.envpol.2024.125437

13. Amaral-Zettler, L. A., Zettler, E. R., & Mincer, T. J. (2020). Ecology of the plastisphere. Nature Reviews Microbiology, 18, 139–151. 10.1038/s41579-019-0308-0

14. Kirstein, I. V., Wichels, A., Gullans, E., Krohne, G., & Gerdts, G. (2019). The plastisphere—Uncovering tightly attached plastic-specific microorganisms. PLOS ONE, 14(4), e0215859. 10.1371/journal.pone.0215859

15. Yu, X., Zhou, Z. C., Shuai, X. Y., Lin, Z. J., Liu, Z., Zhou, J. Y., Lin, Y. H., Zeng, G. S., Ge, Z. Y., & Chen, H. (2023). Microplastics exacerbate co-occurrence and horizontal transfer of antibiotic resistance genes. Journal of Hazardous Materials, 451, 131130. 10.1016/j.jhazmat.2023.131130

16. Zhou, Y., Zhang, G., Zhang, D., Zhu, N., Bo, J., Meng, X., Chen, Y., Qin, Y., Liu, H., & Li, W. (2024). Microplastic biofilms promote the horizontal transfer of antibiotic resistance genes in estuarine environments. Marine Environmental Research, 202, 106777. 10.1016/j.marenvres.2024.106777

17. Liu, Y., Liu, L., Wang, X., Shao, M., Wei, Z., Wang, L., Li, B., Li, C., Luo, X., Li, F., & Zheng, H. (2025). Microplastics enhance the prevalence of antibiotic resistance genes in mariculture sediments by enriching host bacteria and promoting horizontal gene transfer. Eco-Environment & Health, 4(1), 100136. 10.1016/j.eehl.2025.100136

18. Kadac-Czapska, K., Ośko, J., Knez, E., & Grembecka, M. (2024). Microplastics and oxidative stress—Current problems and prospects. Antioxidants, 13(5), 579. 10.3390/antiox13050579

19. Peng, L., Zheng, N., Li, Y., An, Q., Chen, C., Xiu, Z., Li, X., & Wei, Y. (2025). Dynamic evolution of microbial colonization on indoor microplastics: Polymer diversity-driven co-occurrence networks and health risks. Environment International, 207, 109994.

20. Hongjin, C., Ur Rahman, S., Rehman, A., Khan, A. A., & Khalid, M. (2025). Microplastics and antibiotic resistance genes as rising threats: Their interaction represents an urgent environmental concern. Current Research in Microbial Sciences, 9, 100447. 10.1016/j.crmicr.2025.100447

21. Wang, Y. F., Liu, Y. J., Fu, Y. M., Xu, J. Y., Zhang, T. L., Cui, H. L., Qiao, M., Rillig, M. C., Zhu, Y. G., & Zhu, D. (2024). Microplastic diversity increases the abundance of antibiotic resistance genes in soil. Nature Communications, 15(1), 9788. 10.1038/s41467-024-54237-7

22. Wu, C., Song, X., Wang, D., Ma, Y., Ren, X., Hu, H., Shan, Y., Ma, X., Cui, J., & Ma, Y. (2023). Tracking antibiotic resistance genes in microplastic-contaminated soil. Chemosphere, 312(Pt 1), 137235. 10.1016/j.chemosphere.2022.137235

23. Horton, A. A., Walton, A., Spurgeon, D. J., Lahive, E., & Svendsen, C. (2017). Microplastics in freshwater and terrestrial environments: Evaluating the current understanding to identify knowledge gaps and future research priorities. Science of the Total Environment, 586, 127–141. 10.1016/j.scitotenv.2017.01.190

24. Martinez, J. L. (2009). Environmental pollution by antibiotics and antibiotic resistance determinants. Environmental Pollution, 157(11), 2893–2902. 10.1016/j.envpol.2009.05.051

25. Gross, N., Muhvich, J., Ching, C., Gomez, B., Horvath, E., Nahum, Y., & Zaman, M. H. (2025). Effects of microplastic concentration, composition, and size on Escherichia coli biofilm-associated antimicrobial resistance. Applied and Environmental Microbiology, 91, e02282–24. 10.1128/aem.02282-24

26. Wang, H., Xu, K., Wang, J., Feng, C., Chen, Y., Shi, J., Ding, Y., Deng, C., & Liu, X. (2023). Microplastic biofilm: An important microniche that may accelerate the spread of antibiotic resistance genes via natural transformation. Journal of Hazardous Materials, 459, 132085. 10.1016/j.jhazmat.2023.132085

27. Liu, X., Wang, H., Li, L., Deng, C., Chen, Y., Ding, H., & Yu, Z. (2022). Do microplastic biofilms promote the evolution and co-selection of antibiotic and metal resistance genes and their associations with bacterial communities under antibiotic and metal pressures? Journal of Hazardous Materials, 424(Pt A), 127285. 10.1016/j.jhazmat.2021.127285

28. Zheng, Z., Huang, Y., Liu, L., Wang, L., & Tang, J. (2023). Interaction between microplastic biofilm formation and antibiotics: Effect of microplastic biofilm and its driving mechanisms on antibiotic resistance genes. Journal of Hazardous Materials, 459, 132099. 10.1016/j.jhazmat.2023.132099

29. Fajardo, C., Sánchez-Fortún, S., Videira-Quintela, D., Martin, C., Nande, M., D Ors, A., Costa, G., Guillen, F., Montalvo, G., & Martin, M. (2023). Biofilm formation on polyethylene microplastics and their role as transfer vector of emerging organic pollutants. Environmental Science and Pollution Research, 30(35), 84462–84473. 10.1007/s11356-023-28278-2

30. Cholewińska, P., Moniuszko, H., Wojnarowski, K., Pokorny, P., Szeligowska, N., Dobicki, W., Polechoński, R., & Górniak, W. (2022). The occurrence of microplastics and the formation of biofilms by pathogenic and opportunistic bacteria as threats in aquaculture. International Journal of Environmental Research and Public Health, 19(13), 8137. 10.3390/ijerph19138137

31. Loiseau, C., & Sorci, G. (2022). Can microplastics facilitate the emergence of infectious diseases? Science of the Total Environment, 823, 153694. 10.1016/j.scitotenv.2022.153694

32. Maquart, P.-O., Grandjean, L., & Sorci, G. (2022). Plastic pollution and infectious diseases. The Lancet Planetary Health, 6(10), e842–e845. 10.1016/S2542-5196(22)00213-9

33. Nocera, F. P., Attili, A. R., & De Martino, L. (2021). Acinetobacter baumannii: Its clinical significance in human and veterinary medicine. Pathogens, 10(2), 127. 10.3390/pathogens10020127

34. Stoian, I. A., Balas Maftei, B., Florea, C.-E., Rotaru, A., Costin, C. A., Pasare, M. A., Crisan Dabija, R., & Manciuc, C. (2026). Multidrug-resistant Acinetobacter baumannii: Resistance mechanisms, emerging therapies, and prevention—A narrative review. Antibiotics, 15(1), 2. 10.3390/antibiotics15010002

35. Upmanyu, K., Haq, Q. M. R., & Singh, R. (2022). Factors mediating Acinetobacter baumannii biofilm formation: Opportunities for developing therapeutics. Current Research in Microbial Sciences, 3, 100131. 10.1016/j.crmicr.2022.100131

36. Naseef Pathoor, N., Valsa, V., Ganesh, P. S., & Gopal, R. K. (2025). From resistance to treatment: The ongoing struggle with Acinetobacter baumannii. Critical Reviews in Microbiology, 51(6), 1270–1291. 10.1080/1040841X.2025.2497791

37. Sabnis, A., Hagart, K. L., Klöckner, A., Becce, M., Evans, L. E., Furniss, R. C. D., Mavridou, D. A., Murphy, R., Stevens, M. M., Davies, J. C., Larrouy-Maumus, G. J., Clarke, T. B., & Edwards, A. M. (2021). Colistin kills bacteria by targeting lipopolysaccharide in the cytoplasmic membrane. eLife, 10, e65836. 10.7554/eLife.65836

38. Tuon, F. F., Yamada, C. H., de Andrade, A. P., Arend, L. N. V. S., Dos Santos Oliveira, D., & Telles, J. P. (2023). Oral doxycycline to carbapenem-resistant Acinetobacter baumannii infection as a polymyxin-sparing strategy: Results from a retrospective cohort. Brazilian Journal of Microbiology, 54(3), 1795–1802. 10.1007/s42770-023-01015-0

39. Lashinsky, J. N., Henig, O., Pogue, J. M., & Kaye, K. S. (2017). Minocycline for the treatment of multidrug and extensively drug-resistant A. baumannii: A review. Infectious Diseases and Therapy, 6(2), 199–211. 10.1007/s40121-017-0153-2

40. Li, X., Wang, L., Zhang, X. J., Yang, Y., Gong, W. T., Xu, B., Zhu, Y. Q., & Liu, W. (2014). Evaluation of meropenem regimens suppressing emergence of resistance in Acinetobacter baumannii with human simulated exposure in an in vitro intravenous-infusion hollow-fiber infection model. Antimicrobial Agents and Chemotherapy, 58(11), 6773–6781. 10.1128/AAC.03505-14

41. Stabnikova, O., Stabnikov, V., Marinin, A., Klavins, M., Klavins, L., & Vaseashta, A. (2021). Microbial life on the surface of microplastics in natural waters. Applied Sciences, 11(24), 11692. 10.3390/app112411692

42. Nazir, A., Nazir, A., Zuhair, V., Aman, S., Sadiq, S. U. R., Hasan, A. H., Tariq, M., Rehman, L. U., Mustapha, M. J., & Bulimbe, D. B. (2025). The global challenge of antimicrobial resistance: Mechanisms, case studies, and mitigation approaches. Health Science Reports, 8(7), e71077. 10.1002/hsr2.71077

43. Bayot, M. L., & Bragg, B. N. (2025). *Antimicrobial susceptibility testing*. In StatPearls. StatPearls Publishing. https://www.ncbi.nlm.nih.gov/books/NBK539714/

44. Bayot ML, Bragg BN. Antimicrobial Susceptibility Testing. [Updated 2024 May 27]. In: StatPearls [Internet]. Treasure Island (FL): StatPearls Publishing; 2025 Jan-. Available from: https://www.ncbi.nlm.nih.gov/books/NBK539714/

45. O’Toole G. A. (2011). Microtiter dish biofilm formation assay. Journal of visualized experiments : JoVE, (47), 2437. 10.3791/2437

46. Drevets, D. A., Canono, B. P., & Campbell, P. A. (2015). Measurement of bacterial ingestion and killing by macrophages. Current protocols in immunology, 109, 14.6.1–14.6.17. 10.1002/0471142735.im1406s109

47. Dillon, N., Holland, M., Tsunemoto, H., Hancock, B., Cornax, I., Pogliano, J., Sakoulas, G., & Nizet, V. (2019). Surprising synergy of dual translation inhibition vs. *Acinetobacter baumannii* and other multidrug-resistant bacterial pathogens. EBioMedicine, 46, 193–201. 10.1016/j.ebiom.2019.07.041

48. Zhou, X., & Moore, B. B. (2017). Lung Section Staining and Microscopy. Bio-protocol, 7(10), e2286. 10.21769/BioProtoc.2286

